# Using optically-pumped magnetometers to measure magnetoencephalographic signals in the human cerebellum

**DOI:** 10.1101/425447

**Authors:** Chin-Hsuan Lin, Tim M Tierney, Niall Holmes, Elena Boto, James Leggett, Sven Bestmann, Richard Bowtell, Matthew J Brookes, Gareth R Barnes, R Chris Miall

## Abstract

We test the feasibility of an optically pumped magnetometer-magnetoencephalographic (OP-MEG) system for the measurement of human cerebellar activity. This is to our knowledge the first study investigating the human cerebellar electrophysiology using OPMs. As a proof of principle, we use an air-puff stimulus to the eyeball in order to elicit cerebellar activity that is well characterised in non-human models. In three subjects, we observe an evoked component at approx. 50ms post-stimulus, followed by a second component at approx. 85-115 ms post-stimulus. Source inversion localises both components in the cerebellum, while control experiments exclude potential sources elsewhere. We also assess the induced oscillations, with time-frequency decompositions, and identify the source in the occipital lobe, a region expected to be active in our paradigm. We conclude that the OP-MEG technology offers a promising way to advance the understanding of the information processing mechanisms in the human cerebellum.

## Introduction

Our understanding of cerebellar function has undergone a paradigm-shift in recent decades due to the studies of neuroanatomy (Glickstein, Sultan, & Voogd, 2011), neuropsychology (Schmahmann & Sherman, 1998), and functional magnetic resonance imaging (fMRI) (Buckner, 2013; Stoodley & Schmahmann, 2009). Until recently thought of as part of only the motor system, accumulating evidence indicates that the cerebellum is essential for a variety of cognitive and social functions (Sokolov, Miall, & Ivry, 2017). Despite this expanding repertoire of cerebellar functions, there is a marked absence of human electrophysiological studies in this area.

In the domain of magnetoencephalography (MEG), a small body of research has documented cerebellar evoked potentials (Houck et al., 2007; Martin et al., 2006; C. D. Tesche & Karhu, 1997; Claudia D. Tesche & Karhu, 2000), or activity as a part of physiological (Gross et al., 2002; Jerbi et al., 2007; Muthukumaraswamy et al., 2006; Pollok et al., 2004, 2005; Tass et al., 2003) or pathological (Schnitzler et al., 2009; Timmermann et al., 2003) oscillatory networks. These MEG studies provided important insights but they are scarce in comparison with other types of studies, especially fMRI, and have used limited varieties of tasks. It should also be noted that many of them reported cerebellar activation without examining it in detail. This is due to several factors. First, compared to the cerebral cortex, the cerebellar cortex is less favourable to the generation of a measurable MEG signal. Its densely folded anatomy causes a high degree of field attenuation due to locally opposing current sources (Dalal et al., 2013; Hashimoto, Kimura, Tanosaki, Iguchi, & Sekihara, 2003; Tesche & Karhu, 1997). Second, cerebellar neurons are thought to have low firing synchrony based on the small amplitudes observed in local field potential studies (Gerloff, Altenmüller, & Dichgans, 1996). Third, the majority of the cerebellum is located deep in the human cranium, distant from typical SQUID-based MEG sensors. These factors combined make it likely that MEG signals generated in the cerebellum will be smaller and less likely to be distinguishable compared to those produced by the neocortex. Further, cryogenic MEG requires subjects to maintain very still and thus limits the application of many naturalistic movement paradigms relevant to cerebellar dysfunction. The issue is not exclusive to MEG – to date, few studies with scalp cerebellar EEG have been reported (Lascano et al., 2013; Muthukumaraswamy, Johnson, & Hamm, 2003; Todd, Govender, & Colebatch, 2018). It is widely assumed that scalp EEG recordings for the cerebellum suffer from muscle artefacts (Muthukumaraswamy, 2013). Intracranial recordings in the cerebellum are also rare because of their limited clinical indications (Dalal et al., 2013; Niedermeyer, 2004). As a result, the electrophysiology of the human cerebellum is largely understudied compared to the neocortex.

The recent development of optically pumped magnetometers (OPMs) as a tool for MEG provides a new opportunity to investigate cerebellar electrophysiology. OPMs are high sensitivity magnetic field sensors. They do not need cryogenic cooling so can be flexibly placed on the scalp to significantly reduce the sensor-to-brain distance. We have recently built such a wearable OP-MEG system, placing the sensors close to the scalp by mounting them in a 3D printed cast of the head (Boto et al., 2018, 2017). These sensors can be positioned in a dense array over a specific brain region of interest (Tierney et al 2018), although at present only in modest numbers. This system is less susceptible to muscle artefacts compared to EEG (Boto et al., 2018). Moreover, in combination with a field-nulling apparatus (Holmes et al., 2018; Iivanainen, Zetter, Gron, Hakkarainena, & Parkkonen, 2018) one can minimise magnetic field variation at the sensors due to head movement in the ambient field. These characteristics make OP-MEG an ideal candidate for the study of cerebellar electrophysiology, offering possibilities for study of both motor and non-motor tasks.

As a proof-of-concept study, we recorded OP-MEG data when non-noxious air-puffs to the eye trigger blinks. Air-puffs are the unconditioned stimuli (US) in a well-established cerebellar classical conditioning learning paradigm: eyeblink conditioning. In single-unit recording experiments, the stimuli elicit activity in the principal cells of the cerebellum, the Purkinje cells. These cells show in untrained animals (Ohmae & Medina, 2015) both “simple spike” responses driven by input from the brainstem pontine nuclei, as well as “complex spike” responses to climbing fibres that project from the inferior olive to bilateral, predominantly ipsilateral, cerebellar cortex. Animals and humans present comparable behavioural responses to physiological and pathological (cerebellar lesion) interventions in this paradigm (See Freeman & Steinmetz, 2011 for a review of the neural circuitry based on animal studies; Gerwig, Kolb, & Timmann, 2007 for a reivew of human lesion studies). Functional MRI studies in humans also accord well with corresponding electrophysiological studies in animals, with a prominent BOLD response predominantly in the ipsilateral cerebellar cortex (Cheng, Disterhoft, Power, Ellis, & Desmond, 2008; Dimitrova et al., 2002; Thurling et al., 2015). Consequently, cerebellar activation may well be expected in our MEG recording.

We aim here to demonstrate that air-puff driven neural signals in the cerebellum, which have been observed in both the invasive animal and non-invasive human literature (e.g. fMRI), can be measured with OP-MEG. We present evoked and induced responses measured with OP-MEG. Taken together, we show wearable OP-MEG of the cerebellum provides a promising future to examine the cerebellum during human cognition and action, and pathological conditions linked to cerebellar dysfunction.

## Materials and methods

This section is divided into three parts. First, we describe the OP-MEG system. Second, we summarise the experimental procedures for cerebellar activity measurement. Finally, we introduce the inversion scheme used to localise the source activity.

### Participants

Three healthy subjects (1 female, 2 male) aged 27-50, with no history of psychiatric or neurological diseases, participated in the study. All subjects were naïve to the eyeblink conditioning. The research protocol was approved by the University of Nottingham Medical School Research Ethics Committee and the University of Birmingham Research Ethics Committee. Written informed consent was obtained from all participants. The experiments took place at the University of Nottingham.

### OP-MEG System

The OP-MEG system has been previously described in detail (Boto et al., 2018, 2017; Holmes et al., 2018; Tierney et al., 2018). Briefly, the system consists of an OPM sensor array within a customised scanner-cast, and a field-nulling apparatus comprising four reference OPM sensors and field-nulling coils **(Figure 1)**.

**Figure 1.**
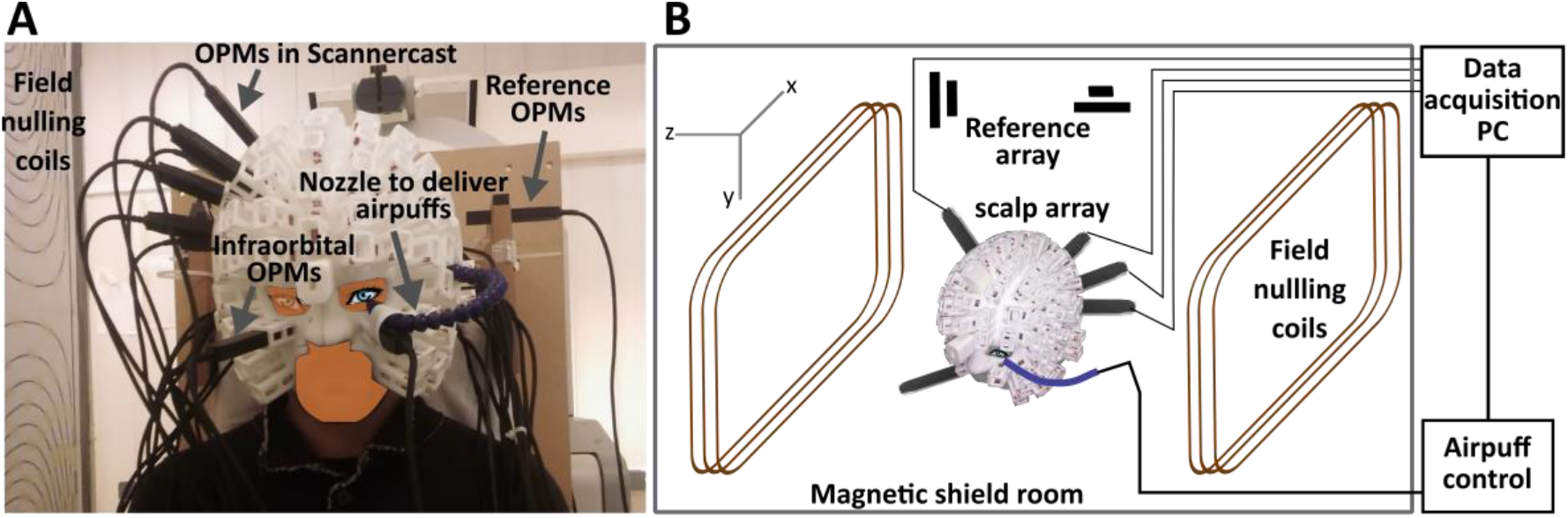
Experimental set-up (A) and schematic illustration (B) of the eyeblink paradigm for cerebellar activity measurement. The participant is seated inside a magnetically shielded room (MSR) wearing a customised scanner-cast. Air-puffs are delivered to the outer canthus of the participant’s left eye via a nozzle. OPM sensors are inserted into slots covering the cerebellum and right somatosensory cortex. Two more sensors are placed in bilateral infra-orbital slots to measure eyeblinks. Field-nulling coils standing either side of the participant carry currents set so as to minimize the residual magnetic field in the MSR, which is measured by 4 reference OPMs prior to scanning. The task-controlling laptop, located outside the MSR, sends synchronizing triggers that were recorded by the data acquisition PC during scanning.

#### Optically-pumped magnetometer (OPM)

The OPM sensors used here (QuSpin Inc., Louisville, CO, USA) have previously been described in detail (Shah & Wakai, 2013; Shah, Osborne, Orton, & Alem, 2018). In brief, each OPM sensor incorporates 3 key components: a 795-nm wavelength laser, a 3 x 3 x 3 mm^3^ cell containing rubidium-87 (^87^Rb) vapour, and a photodiode. The laser spin-polarizes the ^87^Rb atoms in the direction of the laser beam. The photodiode perceives the intensity of the laser transmitting through the cell. At zero magnetic field, the cell is transparent to the laser light and the photodiode detects a maximal signal. A local small field (due to brain activity) causes Larmor procession of the ^87^Rb atoms, which decreases the transparency of the cell because some laser light is absorbed by the atoms. The output of the photodiode shows a zero-field resonance with Lorentzian line shape, allowing the measurement of the field (Dupont-Roc, Haroche, & Cohen-Tannoudji, 1969). In practice, the sensors have a bandwidth of approximately 0-130 Hz with a noise level of ^~^15 fT/√Hz in the 1-100 Hz band, and a dynamic range of ±1.5 nT.

Each OPM sensor also contains a set of three coils which generate three orthogonal fields to cancel out the static field inside the vapour cell up to ^~^50 nT. These on-sensor coils compensate the ambient environmental field in a typical magnetic shield room (MSR) and allow sensors to operate at a fixed position and orientation. To keep OPMs within the dynamic range during naturalistic movement, we further apply an array of bi-planar coils to null the residual field inside the MSR (see the field nulling apparatus section).

#### Field nulling apparatus

The Earth’s residual static field in the Nottingham MSR is ^~^25nT with a maximal gradient of approximately 10 nT/m. This means that even minimal head movements (e.g. a 4 degree rotation) can easily generate signals which exceed the OPM’s dynamic range. In order to mitigate these effects we used a set of bi-planar coils to generate magnetic fields which counteract the remaining static field as measured by four reference sensors close to the head. By applying this method, the dominant component of the static field and field gradient can be diminished by factors of 46 and 13 respectively, in a volume of 40 × 40 × 40 cm^3^ encapsulating the head (Boto et al., 2018; Holmes et al., 2018).

#### Scanner-cast design

Data from an anatomical T1-weighted MRI scan was used to generate a 3D mesh representing the outer surface of each participant’s scalp. This 3D mesh was used in 3D printing to shape the inner surface of a nylon scanner-cast, as described in (Boto et al., 2017; Meyer et al., 2017; Tierney et al., 2018), with sockets around the outer surface to hold the OPM sensors. Importantly, the mesh and subsequently produced wearable helmet provide accurate socket and thus sensor positions and orientations, which are in the same coordinate space as the MRI image. Therefore, no coregistration is needed and the sensor positions are defined as the centre of the socket bases with a 6.5 mm offset, which takes into account the stand-off distance between the outside of the sensor housing and the centre of the vapour cell (Shah, Osborne, Orton, & Alem, 2018). Here, with limited sensors available, we placed the sensors proximal to the cerebellar and somatosensory cortices. **Figure 1** shows the sensor configuration in a typical subject. Across the 3 participants thirteen to nineteen posterior sensors and four to six somatosensory sensors were used. Additionally, two sensors were placed in bilateral infra-orbital slots for eye-blink detection **(Figure 1.A)**.

### Experiment: Eyeblink paradigm

Eyeblinks were elicited by a 32 ms air-puff delivered through a nozzle mounted on the scanner-cast **(Figure 1.A)**; essentially a pressurised air cylinder (1 Bar) fed into a 10 m semi-rigid plastic tube (2mm internal diameter), under the control of a bespoke pneumatic valve controller. Some details of the pneumatic delivery system are given in Leonardelli et al., (2015). The nozzle directed the air-puff to the outer canthus of the left eye from a distance of approximately 2-4 cm, individually adjusted to evoke a visible blink after each delivery, but without discomfort. The arrival time of the air-puff was calibrated off-line using a microphone and was relatively insensitive to distance of the nozzle over a limited range (^~^1 ms/cm). Subjects received four contiguous 12-minute blocks of stimulation. To equate the task to the baseline phase of a previously validated eyeblink conditioning paradigm (Cheng et al., 2008), each block constituted 200 trials: 140 trials of air-puffs, 50 trials of a 550-ms binaural tone (2800 Hz), and 10 paired trials with the tone co-terminated with air-puff delivery; trial order was randomized in sets of 20. A total of 600 air-puff trials were recorded per subject. Every trial began with a random wait of 1-2.5 seconds to avoid habituation to puffs; interstimulus intervals averaged to 3.6 s.

To understand the relationship between neck muscle and OP-MEG activity, two additional blocks, with surface electromyography (EMG) recorded by MEG compatible electrodes available on the CTF275 MEG in the Nottingham MSR, were administered in one subject (subject 3), on a separate day. EMG electrode pairs were placed over bilateral cervical splenius capitis (SPL), the main muscles activated during neck extension, based on (Sommerich, Joines, Hermans, & Moon, 2000).

### OP-MEG data collection

The OP-MEG data were digitized at a sampling rate of 1200 Hz, using a 16-bit digital acquisition (DAQ) system (National Instruments, Austin, TX) controlled by custom-written software in LabVIEW. The air-puff trigger signals were also acquired through the same DAQ system and sent to both the OPM and CTF MEG (during sessions with concurrent EMG) data acquisition PCs to synchronise air-puff triggers, OP-MEG and EMG data.

### Data analysis

All of the data analysis was performed using SPM12 within the MATLAB environment (Release 2014b, Mathworks Inc., Natick, MA).

#### OP-MEG data pre-processing

OP-MEG Data were filtered between 5 and 80 Hz (or 0 and 200 Hz for eyeblink analysis, explained in more detail below) and each trial epoched between -1000 and +1000 ms relative to air-puff onset. Because OPMs are configured as magnetometers (as distinct from gradiometers which are used in many cryogenic MEG systems) they are susceptible to increased environmental interference. We mitigated interference by constructing virtual gradiometers, which linearly regress the signal recorded by the reference array from the signal recorded at the scalp array (Boto et al., 2017; Fife et al., 1999). Thereafter, data were concatenated across blocks and all trials of data were ranked according to signal variance. Trials with variances higher than [median + 3x median absolute deviation] were rejected (Leys, Ley, Klein, Bernard, & Licata, 2013). For the evoked response analysis, the remaining trials (548, 588 and 585 trials with air-puff stimulation, for subject 1, 2, and 3 respectively) were baseline corrected to the mean of the window 50 ms prior to stimulus onset and then averaged.

#### Spectral analysis

For induced spectral power changes, single trial time-frequency (TF) decompositions in sensor space were calculated for each subject using a Morlet wavelet transform (Tallon-Baudry et al 1998) and then averaged across trials and subtracted out the power of the evoked responses. The wavelet transform was calculated for each time-point between -1000 and +1000 ms, with 76 scale bins corresponding to frequencies between 5 and 80 Hz. For each trial and frequency, the mean power of the interval from 50 ms before stimulus onset until stimulus presentation was considered as a baseline level. The power change in each frequency band post-stimulus was expressed as the relative percentage change from the prestimulus baseline.

#### Eyeblink detection

We identified eyeblink responses from the infra-orbital OPM sensors using the following pipeline: We began with data filtered between 0 and 200 Hz. First, a notch filter at 50 Hz was applied to remove power line noise. Second, a high pass filter at 25 Hz and then a low pass filter at 80 Hz were used. Thereafter, we performed full wave rectification and a final low pass filtering at 20 Hz. A blink peak was identified by the highest amplitude in a trial. For data presentation only, we averaged across 10 consecutive trials.

#### EMG data analysis

EMG Data were high-passed filtered at 40 Hz and rectified. Data were then epoched and concatenated as had been performed for the OP-MEG data. Both average waveforms and time-frequency decompositions were examined and compared with OP-MEG data.

#### Comparing the features OP-MEG to eyeblink and EMG data

We examined the trial-by-trial temporal correspondence between activity of OP-MEG and non-neural electrical sources, including eyeblink and muscle potential, using Pearson’s r-values (Yuval-Greenberg at al. 2008). For all trials, we correlated the latencies of OP-MEG peaks with the latencies of blink peaks. For induced responses, the latency of the maximal power of 5-80 Hz OP-MEG time-frequency spectra was computed for each trial and then correlated with peak blink latency. The same relationship between OP-MEG and EMG peaks was also determined. Besides, EMG and OP-MEG traces were sorted by the peak OP-MEG latency to further inspect the connection between these two types of data.

### OP-MEG source localisation

We evaluated the average evoked response using a dipole fit analysis and the induced power change using a beamformer. In both cases the volume conductor model was the Nolte single shell model (Nolte, 2003), implemented in SPM12, using the scalp boundary from the individual T1-weighted MRI.

#### Dipole fitting

We reconstructed sources of the evoked field data for each subject using the SPM implementation of equivalent current dipole fitting (Kiebel, Daunizeau, Phillips, & Friston, 2008). In brief, the inversion scheme assigned initial means and variances of dipole positions and moments (also called ‘priors’). The final dipole locations and moments were optimised using variational Free energy (Friston, Mattout, Trujillo-Barreto, Ashburner, & Penny, 2007; López, Litvak, Espinosa, Friston, & Barnes, 2014; Penny, 2012). Free energy quantifies a model’s ability to explain the same data (by maximising the likelihood, similar to other dipole fitting routines) while penalising excessive optimisation (heuristically speaking, a complexity penalty). Moreover, models in which the sources have different prior locations can be compared using Free energy. This is particularly useful for comparing different anatomical models of the same data (e.g is the source more likely to have arisen from the cerebellum or the neck muscles?)

We specified the initial mean locations of 6 single dipole models based on the literature: (1) right somatosensory cortex (S1, face area), (2) left S1, face area (Nevalainen, Ramstad, Isotalo, Haapanen, & Lauronen, 2006), (3) right cerebellum (in lobule VI), (4) left cerebellum, lobule VI (Cheng et al., 2008), (5) right and (6) left medial parieto-occipital areas. Models 5 and 6 were used because these areas are not only spatially close to the cerebellum but also known to present blink-related activation (Bardouille, Picton, & Ross, 2006; Liu, Hajra, Cheung, & Song, 2017). We also tested if the evoked responses might have originated from non-neural sources (i.e. posterior neck muscles) using two additional initial mean locations at the right and left posterior neck. The standard deviation of each prior dipole location was set to 10 mm. The mean and standard deviation of each prior moment were assigned as 0 and 10 nA·m. To avoid local maxima, 50 iterations, with starting locations and orientations randomly sampled from prior distributions (i.e. the means and standard deviations of prior locations and moments), were carried out for each model in each subject (50 x 6 = 300 iterations in each subject). For each subject we used the model parameters corresponding to the highest free energy value across all iterations. The maximal free energy values across subjects were then averaged.

#### Beamforming

We used the scalar version of a linear constrained minimum variance beamformer algorithm implemented in the DAISS toolbox for SPM (https://github.com/spm/DAiSS) to localise the source of induced spectral power changes. Two different covariance windows were chosen according to observed sensor level responses **(Figure 3)**. We used (1) a covariance window of 5 to 80 Hz and a time window of -50 to +125 ms and contrasted the 75 to 125 ms post-stimulus period with a pre-stimulus period between -50 and 0 ms and (2) a covariance window of 5 to 30 Hz and a time window spanning -400 to +900 ms and contrasted an individually-based 400ms post-stimulus peak power period with a pre-stimulus baseline between - 400 and 0 ms. The regularization rate λ was set to be 0 (i.e. unregularized) (Barnes, Hillebrand, Fawcett, & Singh, 2004). The source orientation was set in the direction of maximal power. The reconstruction grid spacing was 10 mm.

## Results

### Sensor level responses

We looked at the average evoked response to air-puff stimulation on the cerebellar OPM sensors and the sensors positioned over the contra-lateral somatosensory cortex. **Figure 2.A** shows the average evoked response in posterior sensors (left sensors: blue dotted, right sensors: orange solid) for the three subjects. Two main peaks were observed across subjects at around 50-60 ms and 85-115 ms. The field patterns observed at these peaks were qualitatively dipolar **(Figure 2.A** lower panel). There was a slight latency difference (approx. 2 ms on average) between positive and negative going extrema (e.g. Subject 2, **Figure 2.A** upper panel), indicating a more complex source distribution. We also observed two response peaks from sensors close to the primary somatosensory cortex (anterior sensors: green solid, posterior: blue dotted, **Figure 2.B)**. For each subject, the earliest distinct somatosensory evoked response peaked at 40-50 ms post-stimulus, compatible with the p45m response to facial tactile stimuli (Nevalainen et al., 2006).

**Figure 2.**
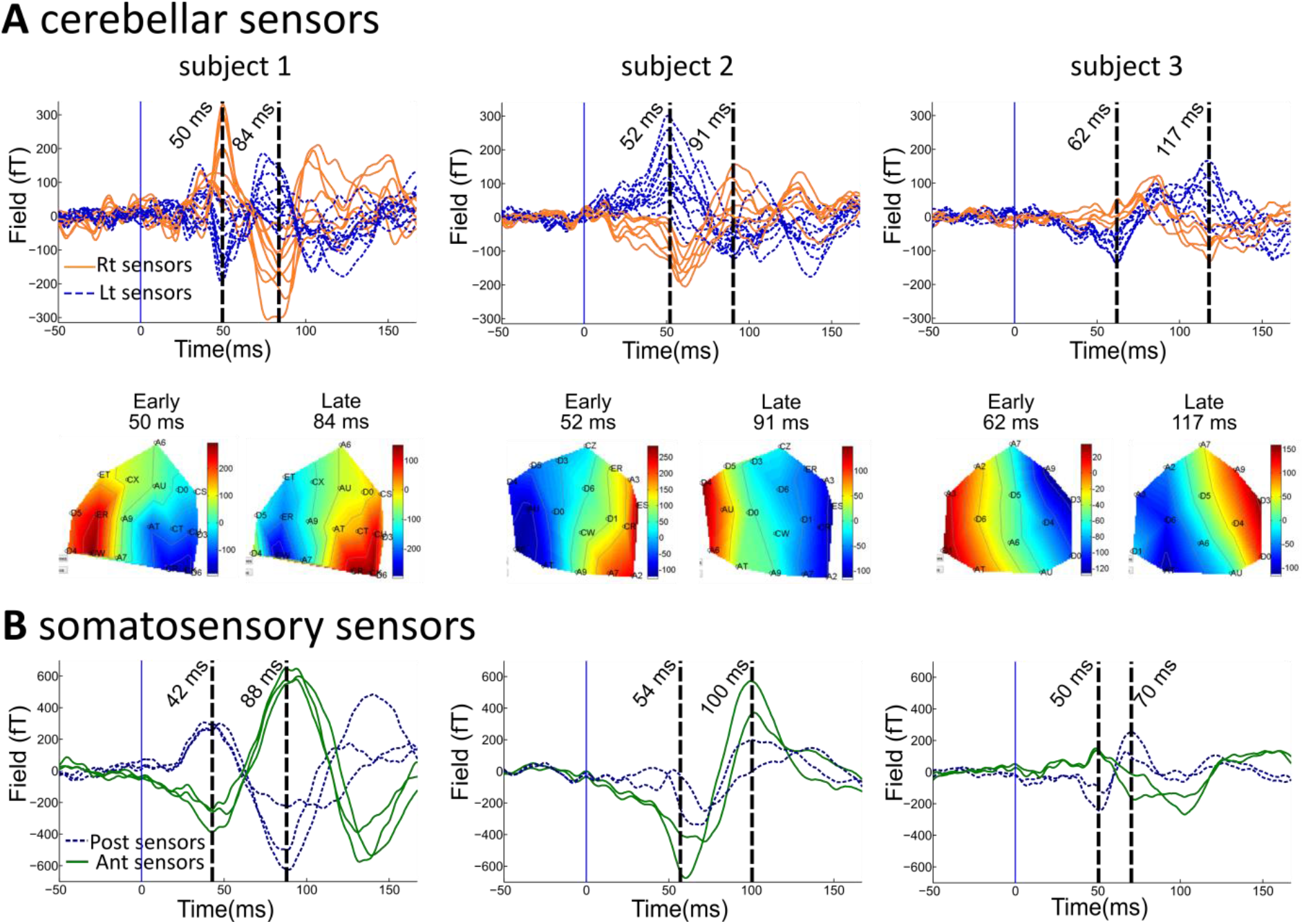
Sensor-level evoked responses following air-puff stimulation. **(A)** Upper panel: air-puff evoked responses over the cerebellar sensor array for 3 subjects. Each trace corresponds to the average signal for one sensor over the posterior cranium, situated left (blue dashed lines) and right (orange solid lines) to the midline respectively. Lower panel: field maps of the evoked field at the latencies of the two distinct peaks; note different sensor layout for each subject. **(B)** Sensor-level evoked responses over the right contra-lateral somatosensory area (anterior and posterior sensors as green-solid and blue dotted curves respectively). Note the time courses are different from the waveforms observed at posterior sensors.

The latency and shape of the blink trace recorded from infra-orbital sensors was dissimilar to the cerebellar evoked responses **(Figure 5.A)**. To test for any contribution to the MEG data from EMG sources in the neck, we sorted OP-MEG trials by the latency of peak EMG. In some sensors, the OP-MEG negative peak seemed to temporally correlate with peak EMG **(Figure 6.B)**. However, peak EMG latency did not time-lock to stimulation and drifted throughout the epoch across trials **(Figure 6.B)**. Besides, these EMG peaks appear to always couple with enhanced negativity of the OP-MEG regardless of the polarity of the evoked response. For example, peak EMG correlated with increased negativity in both OPM channel 11 and 6 **(Figure 6.B)**, while their evoked responses were opposite in polarity **(Figure 6.A)**. Importantly, the EMG-MEG correspondence was only observable outside the time window of evoked responses (before puff stimulation and after 150 ms). We therefore correlated EMG and OP-MEG latencies separately for trials with EMG peaking before and after 150 ms. Only trials peaking after 150 ms had a significant but weak correlation (trials peaking after 150 ms r = .19, p = .01; trials peaking before 150 ms, r = .12, p = .16). While EMG activity peak (at 93 ms) was temporally close to the late peak (at 86 ms) of the OP-MEG evoked responses, no correlation between EMG and OP-MEG amplitude was found at the latency of the late evoked component (r = -.06, p = .30) **(Figure 6.D)**.

We also looked for induced spectral changes in the cerebellar sensors. Figure 3 shows the average time-frequency spectrograms (in percentage change of power from baseline) for the two sensors with the largest power change in each subject. Two types of induced responses can be observed. One was a transient increased broadband power, peaking at ^~^100 ms post puff contact **(Figure 3.A**, left column). The other one was an enhanced activity at ^~^10-30 Hz during 100-900 ms post puff-contact **(Figure 3.A**, right column). The correlation between the latencies of the maximal induced power changes and of peak blinks across trials was only significant in subject 2 (p = 0.03) and was very weak (r = -.09) **(Figure 5.B)**. However, the time-frequency spectra of EMG show a transient enhanced gamma band power at ^~^100 ms **(Figure 6.D)**, which was almost identical to the transient increased broadband response observed in the OP-MEG for this subject.

**Figure 3.**
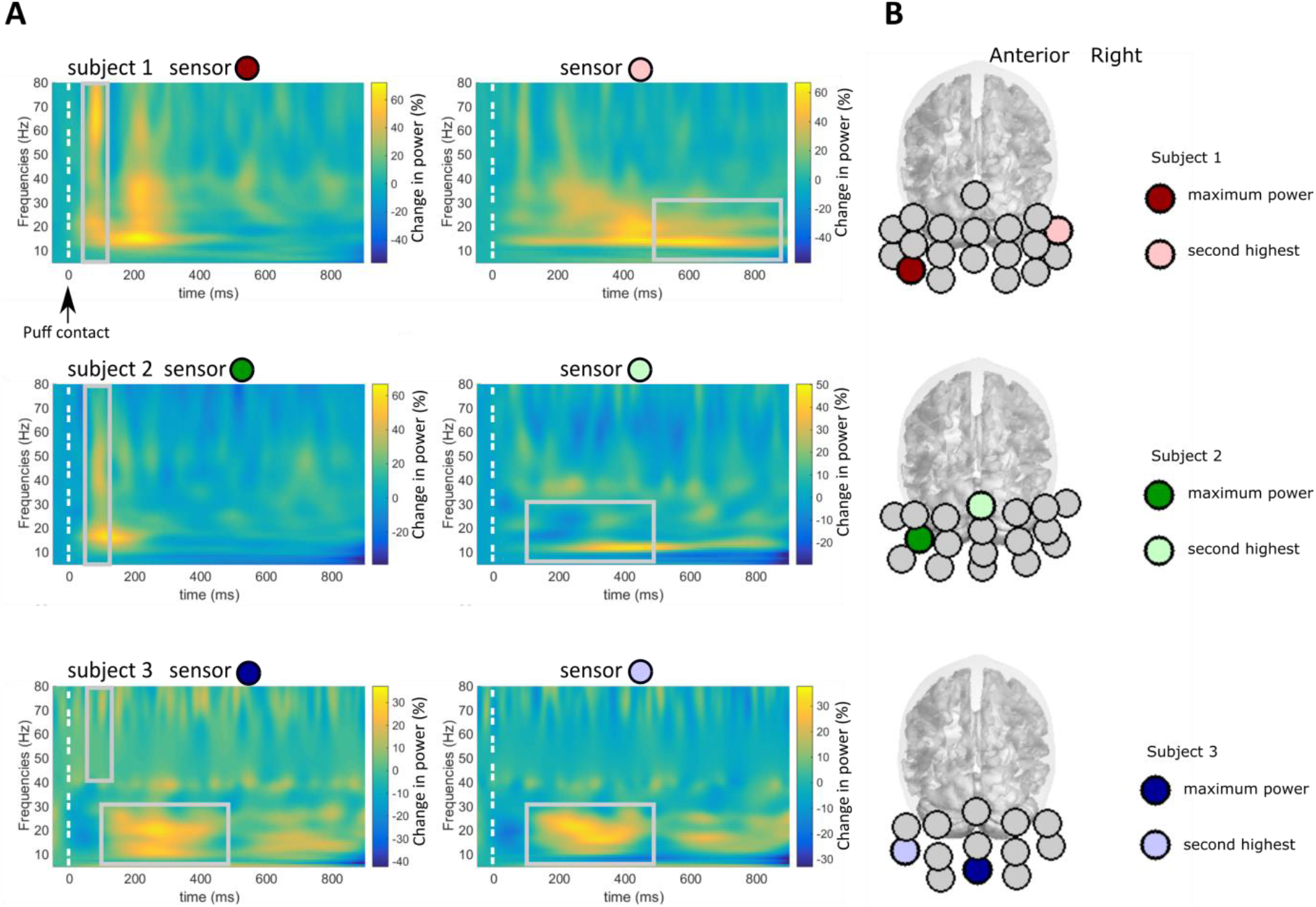
Time-frequency spectrograms showing induced power changes to air-puff stimulation. **A.)** each subject’s average across all trials at the two sensors with the maximal power changes and **B.)** the positions of these sensors are marked as coloured circles for each subject and mapped onto the Montreal Neurological Institute MNI template brain. Two types of enhanced activity can be observed: (1) a transient broadband response peaking at ^~^100 ms (left column) (2) increased power at 10-30 Hz (alpha and beta range) occurring during a 100-900 ms time window with individual latency differences (right column).

### Source reconstruction

**Figure 4** shows the source reconstruction results of early and late evoked responses. **Figure 4.A** compares the free energy values averaged across subjects, for six models initiated at different source locations, for the early (peak latency ±2 timestamp indices, respectively yielding windows of: 48.3-51.7, 50.3-53.7, and 60.3-63.7ms; top panel) and late responses (82.3-85.7, 89.3-92.7, and 115.3-118.7ms; bottom panel). Both cerebellar priors were significantly better than the other models (ΔF > 3; as F approximates the log model evidence, a positive difference of 3 means the preferable model is about twenty times more likely). The left (ipsilateral) cerebellum lobule VI had the highest model evidence for both peaks; although this model was not significantly better than the model of a source in the right cerebellum (ΔF = 0.8 for early and ΔF = 2.2 for late peaks). **Figure 4.B** shows the fitted source locations for subject-specific, early and late time-windows. The source localisations were within the cerebellum or the brainstem/cerebellar peduncle adjacent to the cerebellum. Table 1 shows the MNI coordinates and anatomical labels of each fit.

**Figure 4.**
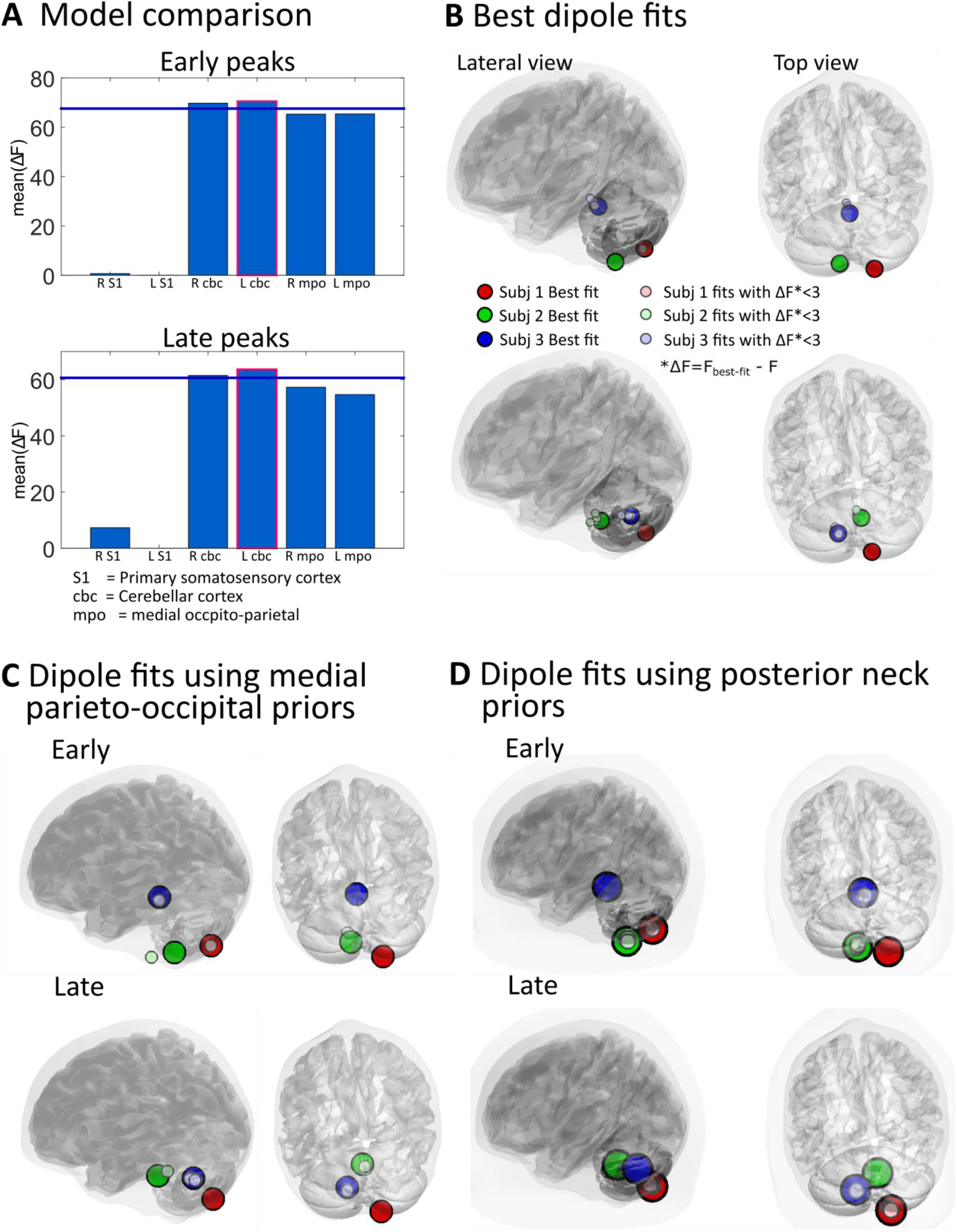
Evoked response source localisation: Single dipole fits using subject-specific early and late peaks of evoked responses. **(A)** Model comparison. Free energy (F) is used to approximate the model evidence of a given source solution. Bars represent the mean Free energy value relative to the poorest model (which was Left S1 both early and late peaks). The left (ipsilateral) cerebellum has the highest model evidence for both early and late peaks. It should also be noted that both the left and right cerebellum are significantly better than the other priors (ΔF > 3, blue line). **(B)** Single dipole fits for each participant. Large circles represent the source locations of models with highest evidence for each individual. Smaller circles are models which are suboptimal but not significantly so (ΔF = F_best-fit_ - F < 3). **(C)** Single dipole fits for each peak and subject when using priors in either left or right medial parieto-occipital area. **(D)** Single dipole fits for each peak and subject using posterior neck priors. As in **(B)**, large circles in both **(C)** and **(D)** represent the best fits for each subjects and small circles represent fits which have ΔF = F_best-fit_ - F < 3. The final estimates of source positions in **(C)** and **(D)** are largely in agreement with the dipole fits using more carefully selected physiological priors in **(B)**.

**Table 1.**
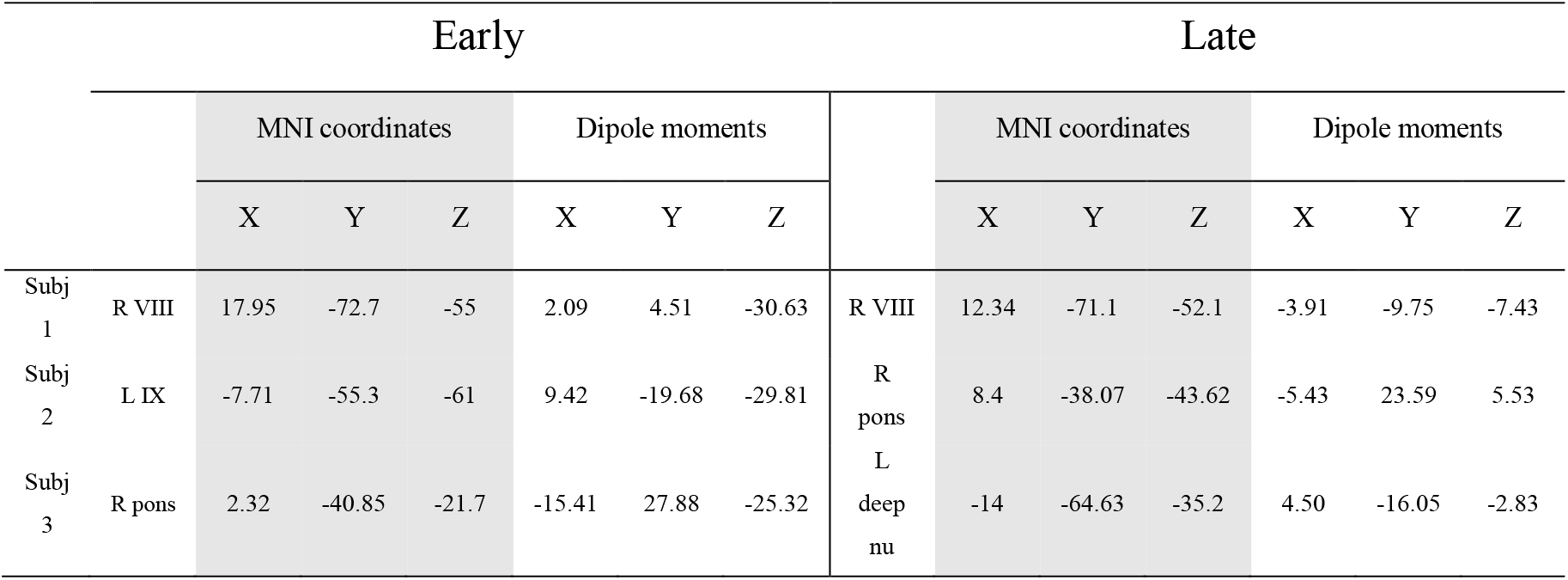
Location of dipole fits for early and late components of the evoked response in 3 subjects. Location labels were based on (Diedrichsen, Balsters, Flavell, Cussans, & Ramnani, 2009).

Dipole locations converged into the cerebellum even when using priors in the medial parieto-occipital areas (model 5 & 6, see **Figure 4.C**). We also undertook additional single dipole fitting using extra-cranial priors at the posterior neck, approximating the SPL muscle, and found similar results **(Figure 4.D)**.

In order to locate the two types of sensor-level induced power changes, we performed the beamforming analysis with two different covariance windows. For the transient broadband response, a window covering a 5-80 Hz frequency band and a time range of -50 to +125 ms was used and we contrasted the power between time windows of +75 to +125 ms and -50 to 0 ms. For the response in alpha and beta range, a window covering a 5-30 Hz frequency band and a time range of -400 to +900 ms was used and we contrasted an individual peak-based 400ms post-stimulus period (500-900 ms for subject 1, 100-500 ms for subject 2 & 3, see **Figure 3.A**) with a pre-stimulus baseline between -400 and 0 ms. For the broadband component, the global maxima were located at the posterior neck. For the induced response in alpha and beta band, the sources were localised in the medial occipital area. Table 2 showes the MNI corrdinates of the global maxima.

**Table 2.** Location of global maxima for broadband and alpha & beta band induced responses in 3 subjects.

|  |  | Broadband |  |  |  |  | Alpha & Beta |  |  |
| --- | --- | --- | --- | --- | --- | --- | --- | --- | --- |
|  |  | MNI coordinates |  |  |  |  | MNI coordinates |  |  |
|  |  | X | Y | Z |  |  | X | Y | Z |
| Subj1 | L neck | -1.25 | -68.34 | -70.23 | R Occipital |  | 0.05 | -90.59 | -7.60 |
| Subj2 | L neck | -5.81 | -41.18 | -63.58 | R Occipital |  | 29.86 | -56.58 | 6.79 |
| Subj 3 | R neck | 12.55 | -88.93 | -63.53 | R Occipital |  | 21.71 | -68.16 | 1.61 |

## Discussion

We have demonstrated that a small OPM array with less than 20 sensors can detect cerebellar evoked responses during unconditioned eyeblinks elicited by brief air puffs. The evoked responses had early (^~^50 ms) and late components (^~^90-110 ms). The source localisation of these evokes response was in the ipsilateral cerebellum, generally consistent with previous neural recordings in animals and fMRI in humans.

In the small number of published MEG studies, cerebellar activation to somatosensory stimuli has been identified by either using *a priori* assumptions of an equivalent current dipole (ECD) in the cerebellum (Tesche & Karhu, 1997; Tesche & Karhu, 2000) or by using a beamforming source reconstruction (Hashimoto et al., 2003). The Hashimoto study (Hashimoto et al., 2003) showed a four-component response in the cerebellum following median nerve electrical stimulation, and the authors made putative assignment of these peaks to different cerebellar inputs. The Hashimoto study showed robust source level images, again lateralized predominantly localized to ipsilateral, medial cerebellum, but was based on 10,000 trials. By contrast, our results are apparent at sensor level and are based on 550-580 trials per participant. We anticipate that a further reduction in required trials could be achieved by bespoke design of a cerebellar scanner-cast, to optimise sensor locations to further increase signals.

Due to the nature of the eyeblink paradigm, there were eye and eyelid movements in almost every trial. We thus examined the relationship between infra-orbital blink signal and OP-MEG data in detail to understand if ocular sources might contribute to the MEG sensor level activity, even though artefacts from the corneo-retinal dipole and extra-orbital muscles have been shown to be highly focal and limited to fronto-central sensors in the MEG (Carl, Açik, König, Engel, & Hipp, 2012; Muthukumaraswamy, 2013). We found that eyeblink occurs relatively late (^~^100-140 ms) after stimulus onset **(Figure 5.A & B)** and, importantly, peaked after the later cerebellar component, for each subject. Blink is therefore an improbable source of these evoked responses. On a trial-by-trial basis, we found no or only a very weak temporal correlation between the latencies of the maximal spectral power changes and blinks **(Figure 5.B)**. Thus, the induced responses were also not likely to be products of blinks.

**Figure 5.**
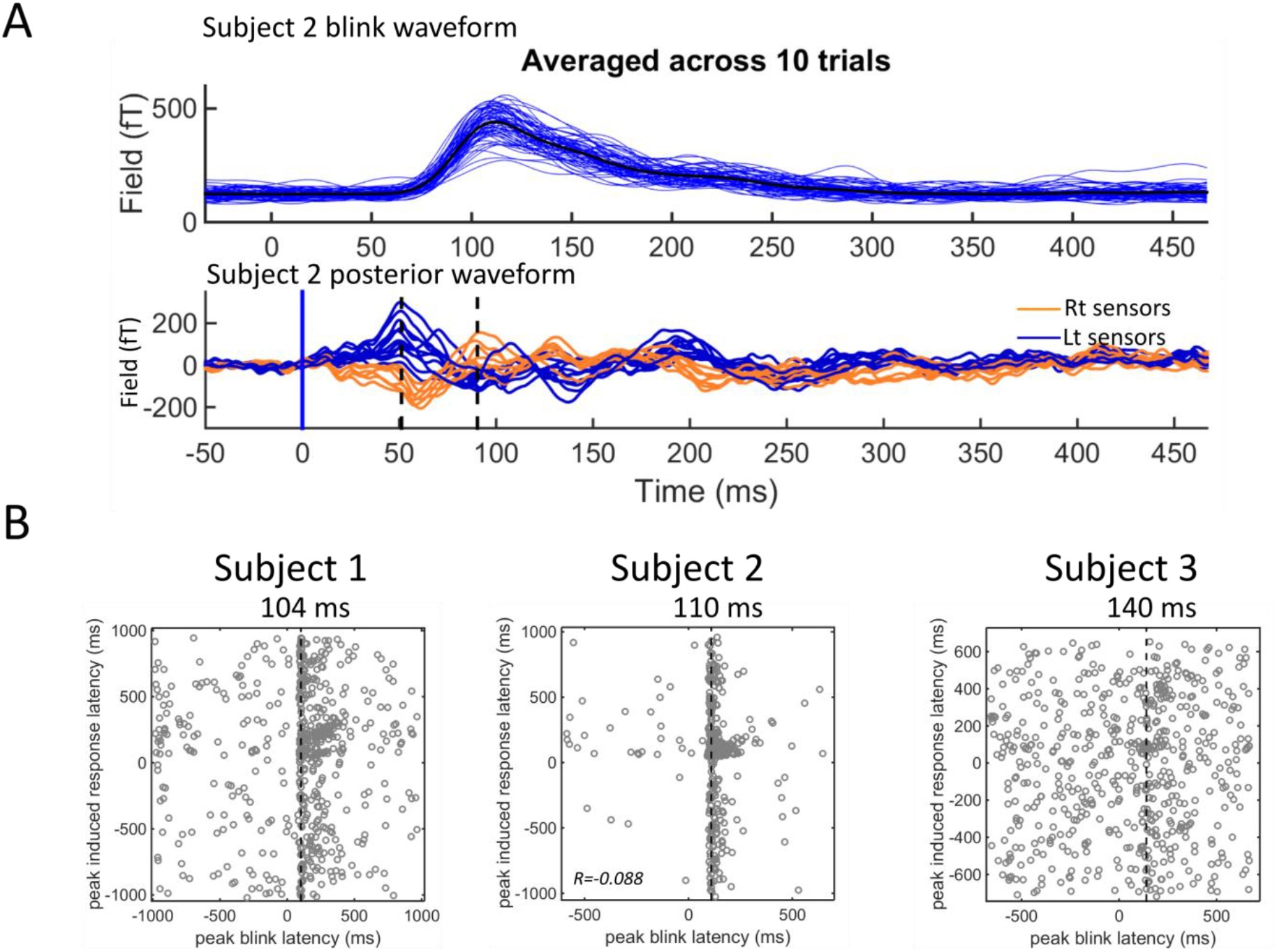
Comparing cerebellar MEG with infra-orbital eyeblink data. ***(A)** Comparison of infra-orbital blink and mean posterior waveforms of subject 2: (Upper) Blue lines are blink traces shown in blocks of 10 trials and the bold black line is the average of all trials. The time course of the blink waveform, constituting of a single peak at 110 ms, is dissimilar to the OP-MEG waveform (as shown in the lower panel). **(B)** The relationship between latencies of maximal induced response and latencies of blink peaks, as shown by the scatter plot for all trials in each subject. For each trial, the latency of the absolute maximal peak within the 5-80 Hz MEG spectrogram data and between - 1000^~^+1000 ms and was determined and plotted against the latency of the blink maximum. Note that these maxima are not necessarily time locked to the air-puff stimulus. Subject 2 showed a significant (p= .03) but extremely small Pearson r value* (r = -.088) (R-squared = .0078; meant that less than 1% of variance is explained the linear model of eyeblink and MEG peak latency). *Correlations were not significant in the other two subjects (subject 1: r = .007, p = .87 and subject 3: r = .053, p = .20). The vertical line of each graph represents the modal latency of the maximum in the blink signal for each subject. Note that they are later than the second MEG peak of each individual (as can be seen in **Figure 2.A**)*.

Another source of non-neural artefact to be considered is the neck muscle because it is nearby the cerebellum and the sudden air puff does lead to some reflex head motion, visible in early acclimatization trials. However, the evoked responses are unlikely to be caused by muscle activity, as dipole fits converged to the cerebellum even when using posterior neck priors **(Figure 4.D.)**. Further, there was no correlation between the amplitude of the peak surface EMG and the OP-MEG response (r = -.06, p = .30) (**Figure 6.C**, right subplot). A weak temporal correlation (Pearson r = .19, p= .01) between EMG and OP-MEG activity broke down during the time window of evoked responses (*r = .12, p = .16;* **Figure 6.B & C**). Regarding the induced responses, the time-frequency spectra of neck muscle EMG **(Figure 6.D)** revealed a transient broadband component at ^~^100 ms and 40-80Hz, whose time and frequency features closely resemble the induced response in the OP-MEG. Beamforming analysis also localised the source of this transient enhancement of gamma band at the posterior neck. We thus conclude that this broadband induced response was likely to be due to muscle artefacts.

**Figure 6.**
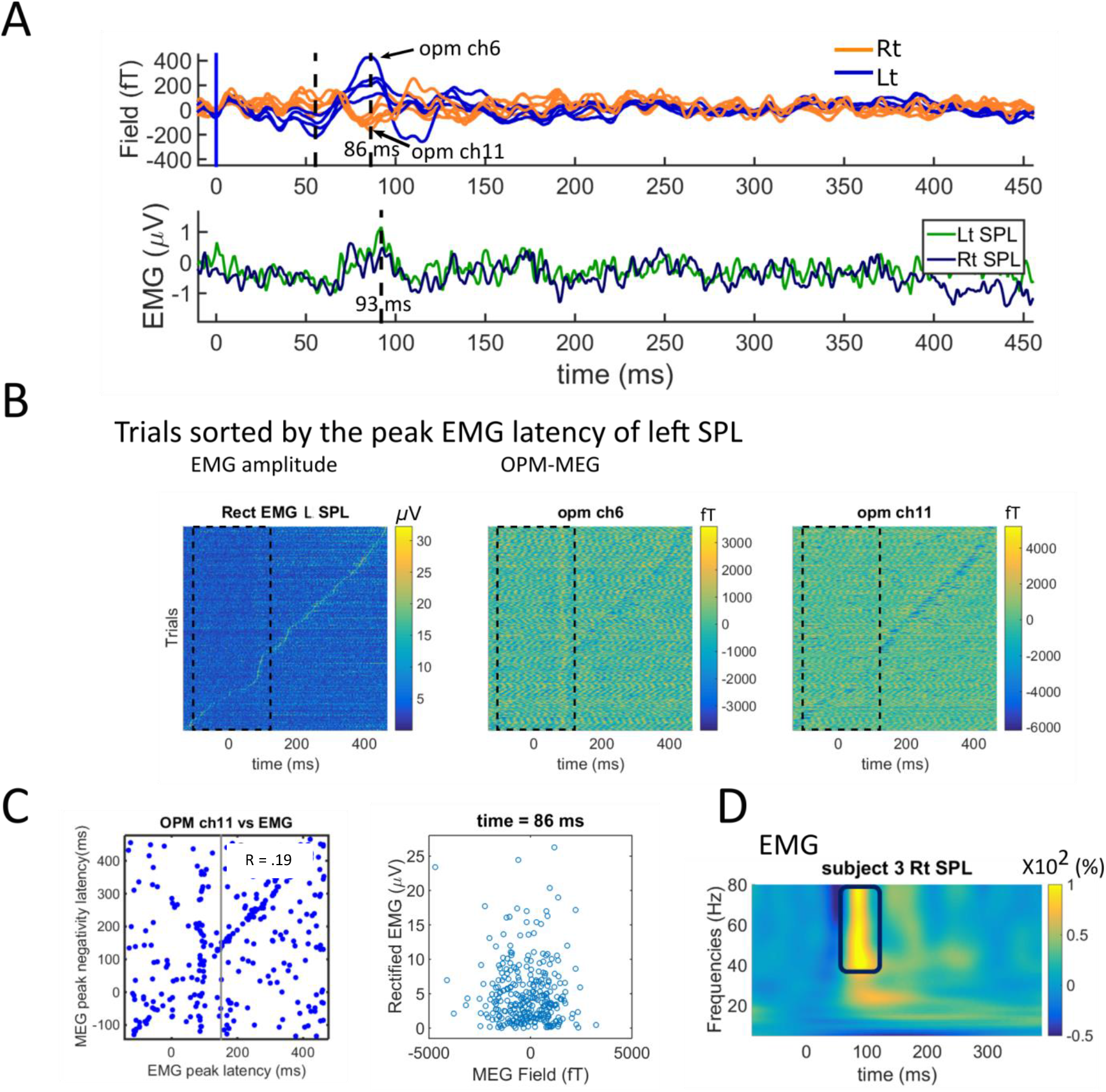
Comparing cerebellar MEG with neck EMG data. **(A)** Comparison of the average posterior waveforms (upper) and rectified EMG (lower) of subject 3: EMG activity peaks (93 ms) later than the evoked responses of OP-MEG, though close to the late peak (86 ms). Note that the OP-MEG evoked responses appear before 150 ms **(B)** EMG and OP-MEG trials sorted by the latency of peak EMG activation, Subject 3. The colour represents the amplitude of EMG and OP-MEG for all trials. Peak EMG latency drifts overtime across trials. Though in some OPM sensors (six sensors out of all. Two are shown here) the MEG negativity peak seems to temporally correlate with the peak EMG, this correspondence is only observable outside the time window of evoked responses. (before puff stimulation and after 150 ms) **(C)** We performed correlation of latencies separately on trials with EMG peaking before and after 150 ms; Subject 3. The Pearson r for trials with EMG peaking before 150 ms (i.e. the time window of evoked responses) is not significant (r = .12, p = .16) and for trials peaking after 150 ms is a weak but significant correlation (r = .19, p = .01). There is no correlation between rectified EMG and OP-MEG amplitude at the time of late MEG peak (r = -.06, p= .30). **(D)** Time-frequency spectrograms of rectified EMG at the splenius capitis (SPL) of subject 3. A transient broadband increased power at gamma range at ^~^100 ms post-stimulation can be seen.

To our knowledge, this is the first MEG study to examine the neural response due to the unconditioned stimulus (US, i.e. air-puffs) of eyeblink conditioning. Being a model system for cerebellar learning, the neural circuits of both unconditioned and conditioned blink responses have been extensively investigated in animals. In untrained rodents, single-unit recording from Purkinje cells in ipsilateral cerebellar cortical lobule VI show that air-puffs elicit climbing fibre inputs, with high probability, peaking around 15-50 ms post-stimulus (Mostofi, Holtzman, Grout, Yeo, & Edgley, 2010; Ohmae & Medina, 2015). Importantly, excitation of the very powerful climbing fibre-Purkinje cell synapse, simultaneously activating the dendrites of a spatially aligned set of Purkinje cells has been considered to be the most probable source for a strong open field detected by MEG recordings (Tesche & Karhu, 1997). According to animal local field potential recording, the US reaches the cerebellum first via an early mossy fibre response and then the aforementioned climbing fibre response (Hesslow, 1994; Mostofi et al., 2010). Our OP-MEG evoked responses showed two peaks, the first tightly clustered around 50-60 ms while the second occurred 80-110 ms post-puff with significant inter-individual differences in latency. These first human data are thus broadly consistent with the animal literature, and the climbing fibre signal might contribute to the early 50 ms response. Previously, similar qualitative features including multiple components (Hashimoto et al., 2003; Tesche & Karhu, 2000) and inter-individual latency variability (Tesche & Karhu, 1997) responding to simple somatosensory input have also been found in human MEG at source levels. Hashimoto et al. speculated that a component at 50-70 ms was driven by the median-nerve induced activation of climbing fibres.

The inversion scheme found the left (ipsilateral) lobule VI was the locus of the best fitting single-dipole model of these evoked responses. This source location is in agreement with previous findings (Cheng et al., 2008). However, it should be noted that in all three subjects, the best evoked-response fits located close to the midline, more medial than lobule VI. Besides, for both early and late peaks of the evoked field, the differences of Free energy values between single dipoles in the right and left cerebellum lobule VI were small (ΔF < 3, **Figure 4.A**). One potential cause of this finding is that although ipsilateral lobule VI is the dominant neuronal source of the US signal, concurrent (but weaker) neuronal activation has been found in the contralateral lobule VI in both human fMRI (Dimitrova et al., 2002; Thurling et al., 2015) and in animal studies (Mostofi et al., 2010), so that a two-dipole model might be more accurate. When fitting evoked fields with two-dipole priors, the majority of best fits were indeed paired bilateral cerebellar sources (figure not shown here). These results support the hypothesis of bilateral activation in the cerebellum by unilateral air-puff stimulation. This said, we admit the current results cannot conclusively locate the sources at a resolution of cerebellar lobules (Schmahmann et al., 1999).

In this proof-of-principle study, we also searched for induced responses to air-puff stimulation. At sensor level, two types of induced responses were observed: a brief increased broadband power at ^~^100 ms and an increased power in a lower-frequency range (10-30 Hz) between 100^~^900 ms. However, as has been mentioned, the source of the former component was found to be outside the skull and likely to be caused by neck muscle activity. While we could not observe any overt head motion, the head was freely supported, with neck muscles activated and it is possible that reflex responses to the air-puff stimulus caused this effect. On the other hand, the alpha and beta band response were located in the medial occipital area. It is possible that the lower-frequency component reflects neural responses to the transient changes of visual input during blinks, as have been reported previously (Bardouille et al., 2006; Liu et al., 2017). However, further studies will be needed to confirm the nature of this induced lower band response.

Our current tests have been limited by the number of OPM sensors available. Given the relatively greater depth of the cerebellum, its complex architecture, and the diffuse spatial signature seen in the evoked responses (see the fieldmaps in the **Figure 2.A.**, lower panel), the source localisation would likely benefit from a denser and spatially extended sensor coverage. This will shortly be available with a next generation OPM sensor. Additionally, it would be useful to design scanner-casts that allow sensors to be placed over the upper neck to provide greater coverage of the inferior cerebellum, which was just below the lower margin covered by the current setup. Finally, to give confidence in the identities and localisations of our findings, we intend to extend the studies to include classical conditioning and extinction of the eyeblink, and thus aim to track the predicted changes in the responses to the unconditioned and conditioned stimuli across the learning and extinction phases.

## Conclusion

We have demonstrated an OP-MEG system that can be used to study the electrophysiology of the human cerebellum. The similarities between human MEG and animal field potential data in this proof-of-principle task offers promise for future studies to advance our understanding of cerebellar function through non-invasive electrophysiology in humans. Possessing sufficient signal detectability and being wearable, and potentially moveable, we expect the OP-MEG system to be a powerful tool in the investigation of cerebellar functions in both motor and cognitive tasks (Boto et al., 2018; Sokolov et al., 2017; Tierney et al., 2018). This OP-MEG system also has the capacity to fill the “white regions” (Niedermeyer, 2004; Schomer & Lopes da Silva, 2010) of the map of clinical electrophysiology in the cerebellum, with impacts on diseases including movement disorders (Bostan & Strick, 2018), mental disorders (Romer et al., 2017), dementia (Fyfe, 2016), to name a few.

## Acknowledgements

CL was funded by a BBSRC research grant (BB/M009645/1) and by Wellcome grant WT212422. This work was also supported by a Wellcome collaborative award to GRB, MJB and RB (203257/Z/16/Z, 203257/B/16/Z). The WCHN is supported by a strategic award from Wellcome (091593/Z/10/Z). RCM was funded by a Royal Society Leverhulme Senior Fellowship and by Wellcome grant WT212422. We thank Uta Noppeney for use of the pneumatic controller to deliver air-puffs and Jose David Lopez for helpful discussion.

## Additional Information Competing Interests

This work was funded by a Wellcome award which involves a collaboration agreement with QuSpin. QuSpin built the sensors used here and advised on the system design and operation, but played no part in the subsequent measurements or data analysis.

